# *In vitro* selection of a microbial consortium predictive of synergistic functioning along multiple ecosystem scales

**DOI:** 10.1101/2020.11.12.379529

**Authors:** Peter Baas, Colin Bell, Lauren Mancini, Melanie Lee, Matthew D. Wallenstein, Richard T. Conant

**Affiliations:** Colorado State University, Natural Resource Ecology Laboratory, Fort Collins, CO, USA; Growcentia Inc., 500 E Vine Dr, Fort Collins, CO, USA; Colorado State University, Soil and Crop Sciences, Fort Collins, CO, USA

## Abstract

Soil microbes form complex interactive networks throughout the soil and plant rhizosphere. These interactions can result in emergent properties for consortia that are not predictable from the phenotypes of constituents in isolation. We used a four-species consortium to assess the capacity of individual microbial species versus different consortia permutations of the four species to contribute to increased P-solubilization using soil incubations and plant growth experiments. We found that as different combinations of bacterial species were assembled into differing consortia, they demonstrated differing abilities to stimulate soil P cycling and plant growth. The combination of all four microbes in the consortia were much more effective at solubilizing P and stimulating plant growth than any of the individual bacterial species alone. This suggests that *in vivo* functionally synergistic soil microbial consortia can be adept at performing specific ecosystem functions *in situ*. Improving our understanding of the mechanisms that facilitate synergistic functioning examined in this study is important for maximizing future food production and agroecosystem sustainability.

## Introduction

Microbial inoculants are increasingly promoted and adopted to enhance crop productivity, soil health, and overall nutrient use efficiency [1]. Given the key role of the phytobiome for soil ecosystem functioning and plant health, microbiome enhancement and engineering seems like a logical intervention in crop production systems. Yet, many challenges lie between microbial discovery in the laboratory and successful implementation in the field [2]. The functional attributes of isolated bacteria and fungi can be routinely assayed in the laboratory. Microbial genomics identifies the potential for specific functions and traits, and can be predictive of phenotype. For some conserved traits, microbial phylogeny alone predicts function [3]. However, the expressed phenotypes of individual isolates often vary with environmental conditions, including pH, nutrient availability, and in response to stress [4]. As a result, single isolates that show promise in the laboratory often prove ineffective when inoculated into field soils [5, 6].

Predictions of *in situ* functioning of isolates are further challenged by the strong effects of interactions among microbial taxa. Deep learning approaches applied to constructed synthetic communities have revealed causality between microbiome composition and host phenotypes in complex systems under nutrient limited or stressed conditions [7]. Herrera Paredes, Gao (7) showed how different synthetic microbial consortia can affect plant gene expression associated with phosphorus starvation. They suggested that the microbe-microbe interactions were likely responsible for microbial community assembly, which, in turn, allowed for emergent interactions with the host plant to occur. This recent work suggests a framework explaining why broadly efficacious microbial inoculants based on isolates are rare and often provide only a limited solution for improving plant productivity and agronomic efficiency in real-world agriculture applications [8].

Functional microbial consortia may be robust alternatives to single isolates for use as beneficial inoculants in agricultural systems. We define a true functional consortium as two or more taxa that demonstrate enhanced function when interacting relative to any individual constituent alone. Many studies have examined the effects of combinations of microbes on crop yield and other outcomes, often compared to single isolates. However, we are not aware of previous studies that have mechanistically attributed consortium effects to synergy among microbial constituents.

The objective of the current study was to assess how the components of a microbial consortium (made up of four bacterial species) contributed to increased P – cycling functionality using *in situ* soil incubation and subsequent plant growth experiments to measure the capacity of individual microbial species along with different permutations/combinations of the representative mixed bacterial species. We hypothesized that a bacterial consortium co-selected for its ability to mobilize P would outperform any single constituent alone. In order to elucidate P-mobilizing synergies among microbial taxa, we studied a four-species bacterial consortia in a commercially available product, MAMMOTH P^®^ [9]. MAMMOTH P^®^ is an organic liquid microbial soil additive that enhances soil P mobilization and plant yield across many crops [10, 11]. This patented technology was developed using a community-level directed selection approach to optimize consortia interactions resulting in phosphorus solubilization. Using this technology, we conducted a series of experiments to compare the performance of individual bacterial species to all consortia combinations. We tested the combinations at three levels of complexity: (1) liquid culture media, (2) soils, and (3) plant-soil mesocosms. We predicted that P-mobilization and plant yield would increase as the number of constituent bacterial species in the functionally selected consortia increased.

## Materials and Methods

### Microbial Culturing

We used a consortium known to mobilize phosphorus (Mammoth P, Growcentia Inc., Fort Collins, CO, USA). Isolate cultures were grown in Criterion™ Nutrient Broth (Hardy Diagnostics Inc., Santa Maria, CA, USA) from a glycerol stock stored at −80°C. Plant extract (2% (w/w) alfalfa extract)) was inoculated with 5% of culture from glycerol stock and allowed to grow for 2.5 days at 25°C which resulted in cell colony forming units (CFU) >10^9^. Next, the isolate cultures were recombined in previously determined relative proportions (Baas, Bell (10); ***Table 1&2***), by adding 200 μL to 800 μL of proprietary restrictive media [9] low in available phosphorus and high in insoluble forms of inorganic P (i.e. FePO_4_ and AlPO_4_). Ortho-P was determined after 1 and 3 h using the ascorbic acid colorimetric method [12] adapted for microtiter plate measurements on a Tecan plate reader. P-mobilization was calculated by dividing the increase in the concentration of ortho-P (nM) by the time incubated (hours).

**Table 1:** Relative proportions of culture volume for the different treatments for the culture and soil experiment. All treatments were conducted for the culture and soil inoculation experiment while a subset of treatments was selected for inoculation in the plant experiment (*)

| Treatment | Proportion (%) |  |  |  |
| --- | --- | --- | --- | --- |
|  | *E. cloacae* | *C. freundii* | *P. putida* | *C. testosteroni* |
| PCiCoE* | 30 | 20 | 40 | 10 |
| PCiE* | 33 | 22 | 44 | 0 |
| PCiCo* | 0 | 29 | 57 | 14 |
| PCoE* | 38 | 0 | 50 | 12 |
| ECiCo* | 50 | 33 | 0 | 17 |
| PCi* | 0 | 33 | 67 | 0 |
| PCiCo | 0 | 40 | 40 | 20 |
| PCoE* | 43 | 0 | 57 | 0 |
| PCo* | 0 | 0 | 80 | 20 |
| CiE* | 60 | 40 | 0 | 0 |
| CiCo* | 0 | 67 | 0 | 33 |
| CoE* | 75 | 0 | 0 | 25 |
| P* | 0 | 0 | 100 | 0 |
| Ci* | 0 | 100 | 0 | 0 |
| E* | 100 | 0 | 0 | 0 |
| Co* | 0 | 0 | 0 | 100 |

**Table 2:**
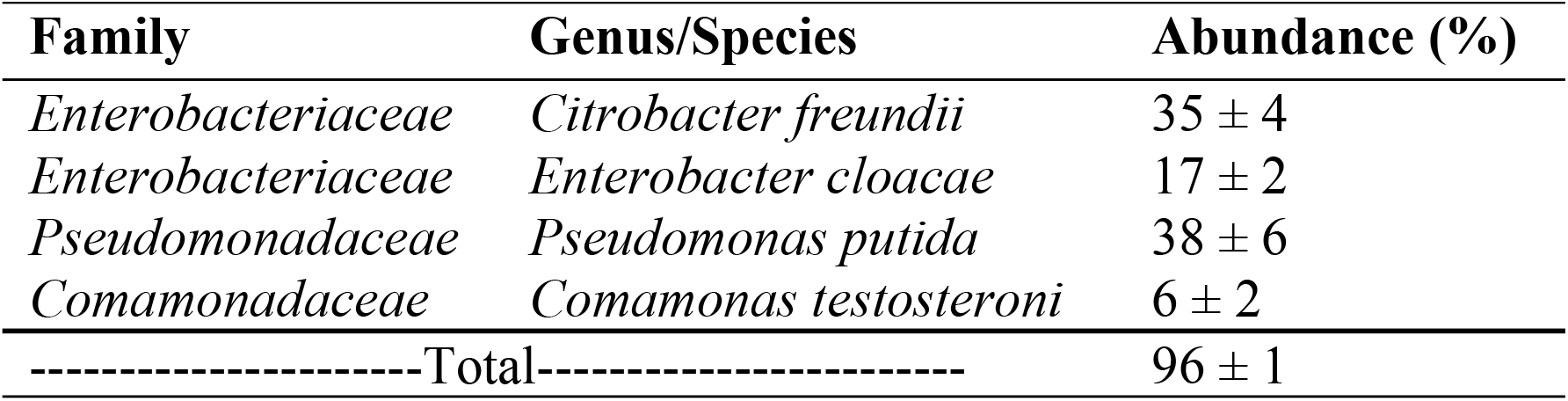
The relative proportions (%) of the top four bacterial species representing >95% of all operationally defined units (out; n=5) in the Mammoth P™ mixture as indicated in Baas, Bell (10).

### Soil Inoculation Trials

Isolates were cultured from a seed bank as described in the previous section. Soil slurries (80g air-dried soil with 400 mL DI water) were prepared from soil sterilized by autoclaving. We used soils collected from the top five cm from three different agricultural systems in Colorado: 1) Agricultural Research Development and Education Center (ARDEC) in Fort Collins, CO, USA; 2) Irrigated corn site at the USDA Central Plains Resources Management Research Station in Akron, CO, USA y (40.15° N 103.15° W) and 3) dryland corn site at the USDA Central Plains Resources Management Research Station in Akron, CO, USA y (40.15° N 103.15° W). The soils were selected for their range in total phosphorus, aluminum and iron concentrations (***Table 3***). The ARDEC soil is of the Nunn soil series (Aridic Argiustolls) and the Akron field site is of the Weld series (Aridic Paleustolls). The initial soil was analyzed for total elemental concentrations of Ca, Mg, Na, K, Ortho-P, total P, Fe, Mn, S and Cl after nitric and perchloric acid extractions using Inductively Coupled Plasma Atomic Emission Spectroscopy (ICP-AES) [13] and carbon and nitrogen concentrations were determined using a total organic carbon (TOC) and total nitrogen (TN) analyzer (Shimadzu Corp., Kyoto, Japan). To estimate the total soil metal (Fe and Al) concentrations as it is related to P-binding capacity we used the binding capacity for Al_2_O_3_ and Fe_2_O_3_ as found in Arias, Da Silva-Carballal (14) to normalize Al concentrations to a Fe_equivalent_ with regards to P-binding capacity. In short, the average QFe and QAl from the Freundlich equation were used to generate a conversion factor for Al (0.18/0.32 = 0.56).

**Table 3:** Soil properties. The numbers indicate the means and associated standard error for the pH. The remaining elements were analyzed on a single composited sample (n=1). Different letters indicate significantly different means (p < 0.05). The elemental concentrations indicate total soil content.

| Soil Type | Location | pH | P (mg kg<sub>soil</sub><sup>-1</sup>) | Fe (g kg<sub>soil</sub><sup>-1</sup>) | Al (g kg<sub>soil</sub><sup>-1</sup>) | Ca (g kg<sub>soil</sub><sup>-1</sup>) | Zn (mg kg<sub>soil</sub><sup>-1</sup>) | Mg (g kg<sub>soil</sub><sup>-1</sup>) |
| --- | --- | --- | --- | --- | --- | --- | --- | --- |
| Corn Dryland | Akron, CO | 5.9 ± 0.1<sup>c</sup> | 199 | 10.3 | 14.4 | 1.7 | 25.0 | 3.3 |
| Corn Irrigated | Akron, CO | 6.8 ± 0.1<sup>b</sup> | 287 | 16.1 | 33.2 | 3.7 | 54.2 | 7.4 |
| Wheat Irrigated | Fort Collins, CO | 8.6 ± 0.0<sup>a</sup> | 444 | 14.3 | 26.1 | 21.0 | 44.7 | 7.8 |

The slurries were well mixed and 450 μL was pipetted into deep-well (2 mL) 96 well plates. The different microbial cultures (***Table 1)*** were added to wells with soil slurry (n=4) and allowed to incubate while gently shaking at 25°C for 8 days. Initial and final available orthophosphate concentrations were determined using 0.5M NaHCO_3_ extractions [15].

### Plant Trials

We used the model plant *Arabidopsis thaliana* whose growth is known to be affected by soil inoculations [16]. *Arabidopsis thaliana* wild type seeds (Edvotek Inc., Washington DC, USA) were planted in four-inch pots with autoclaved soil (50% agricultural soil from the ARDEC (40.39° N 105.00° W) and 50% potting soil (Miracle-Gro® Potting Mix). The soil autoclaving step (which was confirmed effective in removing all native bacteria) was added to allow for the determination of synergistic effects independent of native soil communities. Immediately after planting the seeds, 5 mL of the different cultures outlined in ***Table 1*** were added to each of the tray slots which resulted in previously determined saturation of the root zone. Plant were allowed to germinate for 3-5 days under a mist bench after which they were inoculated a second time with 5 mL and kept moist by daily watering. After two weeks, the plants were moved out of the mist bench and were watered daily in a greenhouse set at 22 ± 2.5°C with a day length of 16 hours. To assess plant growth, we manually measured the number of rosette leaves, the maximum rosette diameter and plant height twice weekly according to Boyes, Zayed (17). Soil available P was determined according to Olsen (18). Plants unable to bolt (emerge) in 32 days which were equally distributed among treatments, were excluded from the analysis.

### Statistics

The differences among treatments in process rates were tested using an analysis of variance approach (ANOVA) and Tukey pairwise comparisons. The relationship between the number of bacterial species and quantified processes was conducted using Spearman’s correlations. All data was checked for normality and, if needed, log or square root transformed to acquire normal distribution. Synergistic or antagonistic effects were determined by subtracting the summed isolate activity from the combination activity and dividing by the summed isolate activity, thus, providing a percentage of the relative non-additive effect. This represents a conservative estimate of non-additive effects since it does not account for reductions in bacterial species-specific abundance when in a consortium. P-mobilization sensitivity was defined as the increase in P-mobilization with the number of bacterial species and abiotic P-mobilization indicates the rate in sterile soils as indicated by the intercept of that relationship. Treatment effect on plant growth metrics were also assessed using repeated measures ANOVA analyses in combination with contrast analyses to determine differences among treatments. All statistics were conducted using JMP 11.

## Results

### Microbial synergy in culture

We found individual bacterial species incubated on low-orthophosphate selective media mobilized P at significantly slower rates than the full consortium (***Figure 1; Table S1***), indicating strong synergistic effects (***Table 4***). The most successful individual species was *C. testosteroni* followed by *C. freundii*, both of which were significantly greater than *E. cloacae* but not than *P. putida.* The simpler consortia with similar mobilization rates to the full consortia (*PCo*, *PE*, *PCiE*. *PCi* and *CiE*) were not significantly different from the full consortia but also not from any other treatments except for the control and the *E* and *P* single strain inoculation. P-mobilization rates within consortia of 2-3 bacterial species were up to 4-fold greater than those of individual bacterial species.

**Table 4:**
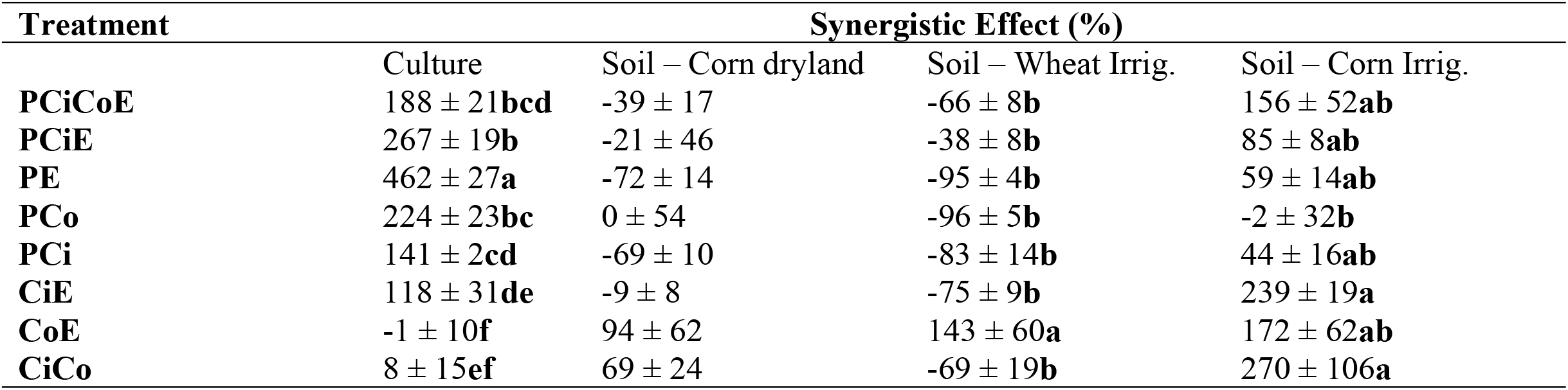
Synergistic effects for a selection of the tested consortia compositions for P-mobilization in culture and three different soils. C = *Citrobacter freundii*; E = *Enterobacter cloacae*; P = *Pseudomonas putida*; Coma = *Comamonas testosteroni*. For each treatment n=4. Irrig. = irrigated. The error represents the standard error of the mean and different letters indicate significant differences among treatments.

| Treatment | Synergistic Effect (%) |  |  |  |
| --- | --- | --- | --- | --- |
|  | Culture | Soil – Corn dryland | Soil – Wheat Irrig. | Soil – Corn Irrig. |
| **PCiCoE** | 188 ± 21**bcd** | -39 ± 17 | -66 ± 8**b** | 156 ± 52**ab** |
| **PCiE** | 267 ± 19**b** | -21 ± 46 | -38 ± 8**b** | 85 ± 8**ab** |
| **PE** | 462 ± 27**a** | -72 ± 14 | -95 ± 4**b** | 59 ± 14**ab** |
| **PCo** | 224 ± 23**bc** | 0 ± 54 | -96 ± 5**b** | -2 ± 32**b** |
| **PCi** | 141 ± 2**cd** | -69 ± 10 | -83 ± 14**b** | 44 ± 16**ab** |
| **CiE** | 118 ± 31**de** | -9 ± 8 | -75 ± 9**b** | 239 ± 19**a** |
| **CoE** | -1 ± 10**f** | 94 ± 62 | 143 ± 60**a** | 172 ± 62**ab** |
| **CiCo** | 8 ± 15**ef** | 69 ± 24 | -69 ± 19**b** | 270 ± 106**a** |

**Figure 1:**
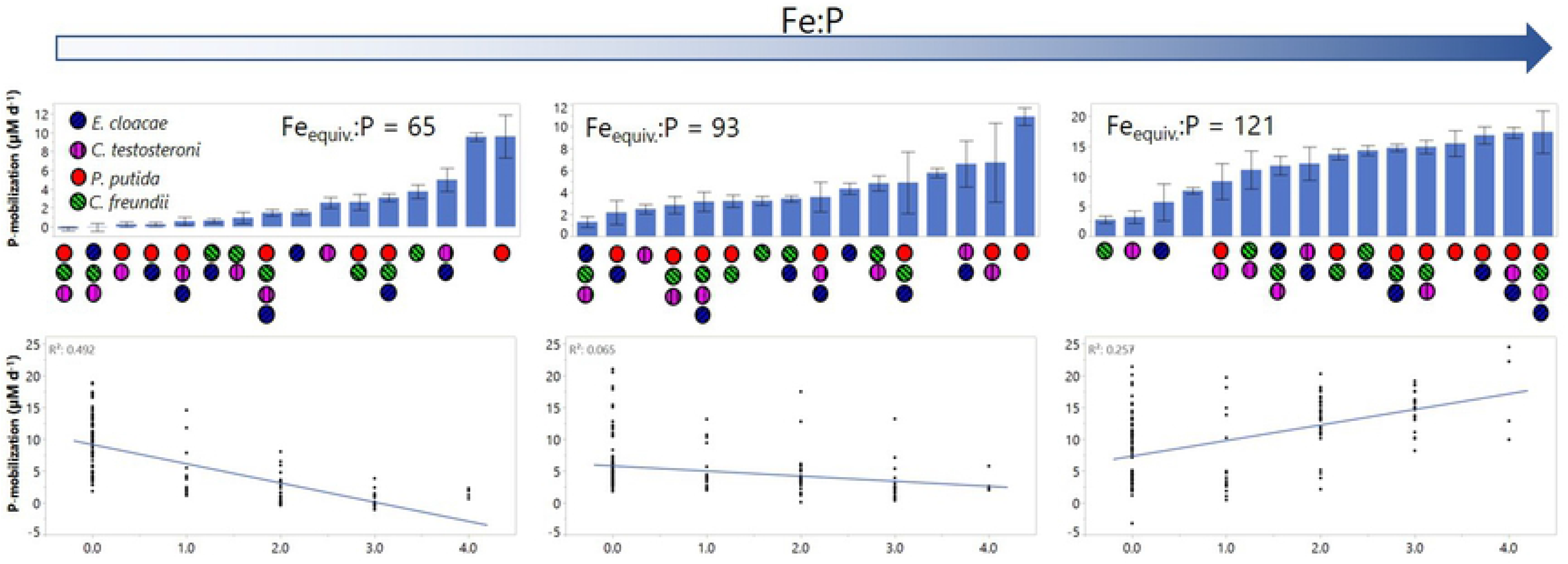
Culture P-mobilization in P-limiting media. The regression on the bottom was significant with p < 0.001. The points and bars represent the mean and the error bars indicate the standard error of the mean. Different letters indicate significant differences. The color and pattern indicate the presence of different combinations of species. For each treatment n=4. Purple/hatched vertically = *C. testosteroni*; Blue/hatched 30° = *E. cloacae*; Green/hatched 330° = *C. freundii*; Red/solid = *P. putida*.

### Microbial synergy in soil

We tested the potential for microbial P-solubilization in soils across an estimated gradient of phosphate saturation of the Fe and Al oxides (Fe_equivalent_:PO_4_). The mobilization of P in soils varied with both inoculum composition and soil characteristics (***Figure 2; Table S2***). In the soil with high a Fe_equivalent_:PO_4_, we found the full consortia, *PE. PCo*, *PE* and *PCiCo* to result in greater P-mobilization rates than the control. In contrast, the soil with low Fe_equivalent_:PO_4_ showed all treatments except for *P*, *Ci* and *CiCo* to result in significantly lower P-mobilization rates. In the soil of medium Fe_equivalent_:PO_4_ no inoculation type resulted in greater P-mobilization than in the control. *PCiCo* and *ECiCo* were even significantly lower in their soil mobilization rate than the control. We found that resource availability, as indicated by the ratio between total P and the total P-binding capacity of Al and Fe (Fe_equivalent_), influenced whether consortium members acted synergistically or antagonistically (***Table 4***). In soils with a greater concentration of potentially bound P (narrow ratio of Fe_equivelent_ to P ratio), the four microbes exhibited an antagonistic relationship, while soils with a wider ratio showed a synergistic response (***Figure 2; Table S2***). Soil P-mobilization sensitivity, defined by the rate at which a greater number of bacterial species increases P-mobilization, was positively correlated to the total Fe_equivalent_:P (***Table 3***; r^2^ = 0.99, *P* < 0.05).

**Figure 2:**
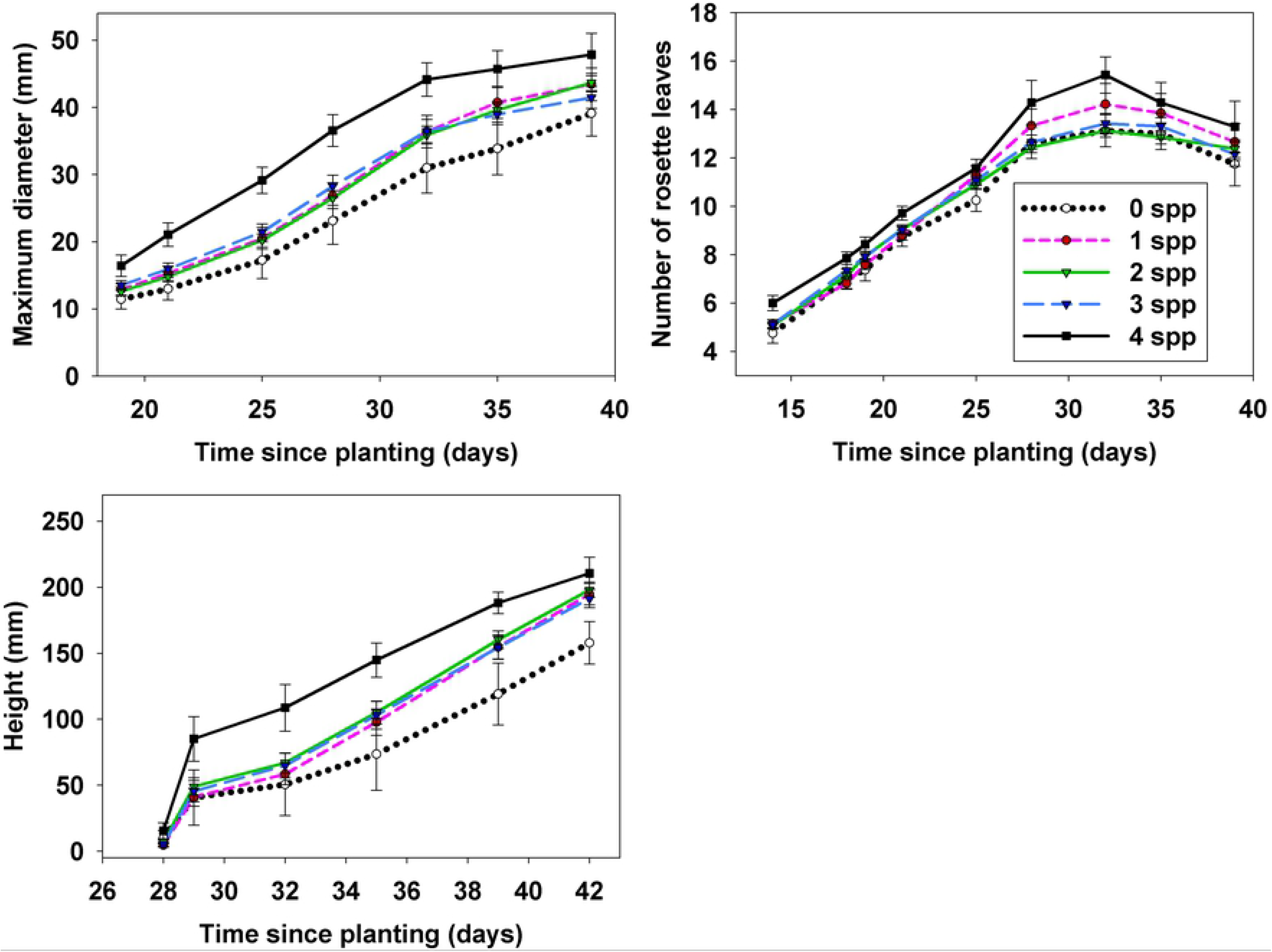
Soil P-mobilization from ARDEC (a), dryland Akron (b) and irrigated Akron (c) soil with different microbial inoculation treatments. The regressions on the bottom for ARDEC (d), dryland Akron (e) and irrigated Akron (f) were significant with p < 0.01. The Fe:P ratio indicates the total Fe-equivalent concentration (including Fe and Al) to total P. The greater the ratio the greater the amount of phosphorus is being bound by Fe and Al oxides. For each treatment n=4. Purple/hatched vertically = *C. testosteroni*; Blue/hatched 30° = *E. cloacae*; Green/hatched 330° = *C. freundii*; Red/solid = *P. putida*. The points and bars represent the mean and the error bars indicate the standard error of the mean. Different letters indicate significant differences.

### Plant Growth and Development

We found that plant development varied widely among inoculum treatments. A repeated measures analysis (29-35 days; p = 0.12) on plant height suggested that the full four species consortia treated plant treatment was greater than the control treatment (p < 0.05). We could not detect specific treatment effects for any of the treatments in the maximum rosette diameter or the number of rosette leaves using repeated measures analysis. We found that increasing the number of bacterial species in the consortium from one to four species increased plant height by up to 32% (***Figure 3***). Further, inocula containing 1-3 bacterial species were greater in height than the control. Residual available P in the soil after plant harvest was greatest in the zero-bacterial species control (44 ± 2 mg kg^-1^) and *CiE* (46 ± 4 mg kg^-1^) treatment and lowest in the *C* (25 ± 5 mg kg^-1^) and four species consortia (28 ± 4 mg kg^-1^) treatments. Inoculation with all four bacterial species reduced the available P by 36% in comparison with the non-inoculated treatment.

**Figure 3:**
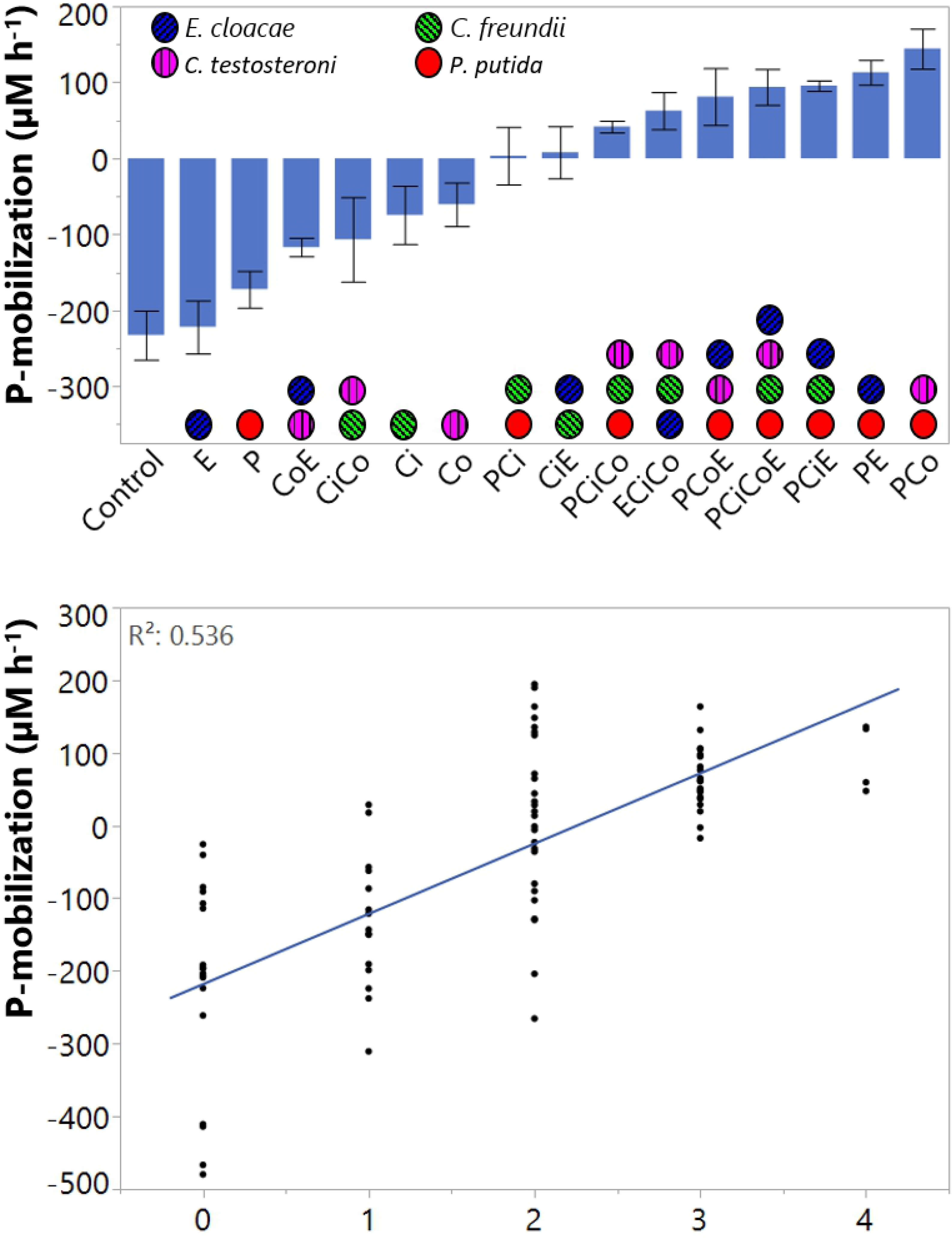
Arabidopsis growth metrics with different numbers of bacterial species (*P. putida, E. cloacae, C. freundii and C. testosteroni*) added. The points represent the mean and the error bars indicate the standard error of the mean. For every treatment n=12.

## Discussion

Across scales–from microplates, soil incubations, and potted plants – we observed a trend of increasing P solubilization and associated benefits with increasing number of taxa within the consortia. Although, in soil, the magnitude and direction of synergistic effects depended on the relative amount of PO_4_ compared to Fe and Al concentrations. At each scale, the performance of the full consortium exceeded that of the best performing individual isolate.

Synergistic microbial interactions have been observed in ecological [6, 19], biofilm [20–22] and bioengineering studies [20, 23–25]. However, net positive interactions appear rare while competition is prevalent [26, 27]. The majority of positive effects are likely carried out by a very small proportion of the microbial community that are strongly metabolically linked. Thus, a small consortium selected for a specific trait may represent a small subset of positive interactions in the overall microbial community. The individual species in the consortium tested in this study rarely exhibited positive effects of the tested trait relative to the control (i.e. P-solubilization, P-mobilization or plant growth). However, the relationships in culture clearly showed that with increasing consortia complexity, the cultures shifted from P-immobilization to P-solubilization.

In synergistic consortia, individual constituents can enhance overall performance through interactive rather than direct effects. Previous research has identified *Comamonas spp*. to be important for P-solubilization [28], yet, the *Comamonas testosteroni* in our study proved important for consortium performance in culture but showed no P-solubilization when inoculated as a single strain. We found the communities with two species to be very inconsistent in P-solubilization rates while communities of 3-4 species consistently solubilized P in culture. These results contradict a theoretical model predicting that synergistic interactions should emerge in consortia greater than three bacterial species [29], but it is not uncommon for synergistic effects to occur within simple ecosystems akin to our laboratory conditions [20, 23–25]. Overall, these findings suggest that although variance was substantial, consortia are often superior to individual strains of the studied four bacteria and have emergent properties with regards to P-solubilization in culture.

The effects of different constructed consortia on P-mobilization were inconsistent across different soils. *Pseudomonas putida* consistently mobilize P independent of soil type, while the direction and magnitude of the synergistic response to increasing consortia size depended on soil type. Surprisingly, even though one of our source soils was alkaline (dominated by calcium oxides), the ratio between Fe_equivalent_ to P appeared to control the sensitivity of P-mobilization, not the combination of Ca, Fe and Al as might be expected since these three elements are part of the dominant three oxides in soil responsible for P-immobilization [30]. This suggests that solubilization of phosphate sorbed to Fe and Al oxides are the main mechanisms used by the selected bacteria to increase available phosphorus. The patterns of antagonistic and synergistic effects follow the principles of stoichiometric theory where microbes mobilize P when bound-P is high (low Fe_equivalent_:P) but immobilize P at low availability (high Fe_equivalent_:P) [31]. Indeed, we found antagonistic effects under a low Fe_equivalent_:P ratio and high synergistic effects under a high Fe_equivalent_:P ratio. This suggests that the effects observed in culture only translate to the soil environment in soils high in occluded phosphate.

In contrast to the results of the soil incubations, when plants were grown in a mixture containing autoclaved soil from the site high in total P concentrations we observed strong synergistic effects with inoculation of increasing consortium size. As would be expected, it is highly unlikely the control treatment remained sterile throughout the experiment but it prevented native microbial interaction from immediately outcompeting the inoculum species. Although, these results do not guarantee these synergistic relationships will also manifest with inoculations in more complicated ecosystems, previous research on the same consortium has found positive effects on plant growth when inoculated in native soil [10]. Microbial P-solubilization has been linked with plant performance in many previous studies summarized by Richardson and Simpson (32) and, similarly, our greenhouse experiment showed that the soils of plants inoculated with larger consortia (which had greater P-solubilization in culture) resulted in increased P-mining. Why did we not observe this effect in the soil incubation? Perhaps plant root exudation shifted the soil stoichiometry to a degree that the inoculated bacteria initiate P-mining [33] which was not present in the soil only incubations. Alternatively, the observed growth enhancement could be the results of wide range of plant-microbial dynamics related to for example indoleacetic acid (IAA), siderophore activities and ACC deaminase [34]. Unlike in the culture and soil experiments, the greenhouse experiment showed strong alignment with the theoretical model by Guo and Boedicker (29) suggesting that a more complex ecosystem including plants is more sensitive to emergent properties from bacteria-bacteria interactions. Inconsistent effects of single-bacterial species microbial inoculants might be explained by the lack of breadth in functional traits [35, 36]. Further, the reduced variance for larger consortia indicates that consortia size controlled not only the enhancement in growth, but enabled plants to more consistently approach their genetic growth potential.

This series of experiments demonstrates that microbial functional traits can emerge within bacterial consortia that are not apparent at the individual taxa level – *i.e.* the whole is more than the sum of its parts. Our study shows that screening individual bacterial species may not yield the same effects when compared to a microbial consortium. Microbial consortia can have superior function compared to individual bacterial species and these synergies may transcend ecosystem scales. Future studies need to be focus on determining the prevalence of these mechanisms in more complicated ecosystems. This work has important implications for efforts to harness the soil microbiome to enhance food production and agroecosystem sustainability.

## Acknowledgements

The authors acknowledge the support of the U.S. Department of Agriculture (USDA) (#2015-67030-22990), Colorado State University (#14BGF19A), the National Science Foundation Directorate for Biological Sciences (DEB-1020540) and Colorado Office of Economic Development and Intl Trade (#19KO BSGR ECFCP7122).

## Supporting information

***Table S1***: P-mobilization rates in culture (nM/h)

***Table S2***: P-mobilization rates in soil (μM/d)

